# Marmoset Anterior Cingulate Area 32 Neurons Exhibit Responses to Presented and Produced Calls During Naturalistic Vocal Communication

**DOI:** 10.1101/2025.02.25.640138

**Authors:** Kevin D. Johnston, Rebekah E. Gilliland, Raymond K. Wong, Stefan Everling

## Abstract

Vocal communication is a complex social behavior that entails the integration of auditory perception and vocal production. Both anatomical and functional evidence have implicated the anterior cingulate cortex (ACC), including area 32, in these processes, but the dynamics of neural responses in area 32 during naturalistic vocal interactions remain poorly understood. Here, we addressed this by recording the activity of single area 32 neurons using chronically implanted ultra high density Neuropixels probes in freely moving common marmosets (*Callithrix jacchus*) engaged in an antiphonal calling paradigm in which they exchanged long-distance “phee” calls with a virtual conspecific. We found that many neurons exhibited complex modulations in discharge rates in response to presented calls, prior to and following self-generated calls, and during the interval between presented and produced vocalizations. These findings are consistent with the conceptualization of area 32 as an audiovocal interface integrating auditory information, cognitive processes, and motor outputs in the service of vocal communication.

**Significance Statement:** Vocal communication is indispensable in the daily lives of social animals including primates. This sophisticated ability requires processing and production of vocalizations across fluid social contexts. Vocal behavior is controlled by a large network of brain areas. Area 32 within the anterior cingulate cortex may be a linchpin of this network, as it is interconnected with both auditory cortical areas and subcortical structures engaging vocal control. This position is ideal for integrating auditory, motor, and cognitive signals serving vocal behaviour. We show for the first time that neural correlates of these three signal types are present in area 32 neurons recorded in freely moving marmosets during naturalistic vocal exchanges. We conclude that area 32 exhibits the properties of an audiovocal interface.

## Introduction

Vocal communication is a complex behaviour that entails interactions between a sender, an auditory signal, and a receiver, all of which operate under the influence of fluidly changing contexts (Bradbury and Vehrencamp, 2000). In primates, this complexity is reflected in the fact that the neural circuits instantiating the linked processes of auditory perception and vocal production of social signals range broadly across the neuroaxis, involving subcortical structures and cortical areas including the anterior cingulate cortex (ACC) (Jürgens, 2009; Grijseels et al., 2023).

A large body of evidence supports a general role of the ACC in the integration of sensory, motor, and cognitive processes (Paus, 2001; Kolling et al., 2016; Amodio and Frith, 2006), extending to those underpinning social interactions (Hadland et al., 2003; Rudebeck et al., 2006). This includes critical aspects of vocal communication such as the auditory processing of conspecific vocalizations and vocal production. ACC areas 32 and 10m/32v are interconnected extensively with auditory cortices including higher order areas responsive to conspecific vocalizations (Tian et al., 2001; Medalla et al., 2007; Medalla and Barbas, 2014) and indeed both ultra high-field fMRI (Jafari et al., 2023; Dureux et al., 2024) and electrophysiological (Gilliland et al., 2024) investigations have revealed robust responses to these stimuli within area 32. Pregenual and subgenual ACC, which encompass area 32, have also been associated with vocal production. These areas send substantial anatomical projections to the periaqueductal gray (PAG) (Müller-Preuss and Jürgens, 1976; Mantyh, 1983), a midbrain region critically involved in vocal control (Jürgens, 1994), and lesions or microstimulation of these areas impair (Sutton et al., 1974) or evoke (Jürgens and Ploog, 1970) vocalizations, respectively. Altogether, the combined weight of anatomical and functional evidence suggests that area 32 has properties consistent with those of an “audiovocal interface” linking perceived and produced vocalizations (Jürgens, 2009).

Beyond the studies noted above, little is known regarding the specific role of ACC area 32 in vocal communication either in or outside of a social context. This is due in part to the relatively limited vocal repertoire of rhesus macaques, and the technical difficulties inherent in conducting electrophysiological studies in freely moving animals with naturalistic tasks that encourage the production of species-typical vocal exchanges. The common marmoset (*Callithrix jacchus*) is a model species within which these challenges can be addressed. As an arboreal species, this small New World primate relies on vocal communication to facilitate social cohesion and survival in the tree canopy (Bezerra and Souto, 2008). One aspect of this communication is the species-typical long-distance “Phee” call, which is used to maintain contact between group members and typically expressed in a call-and-response pattern (Takahashi et al., 2013). These antiphonal exchanges are reliably produced in the laboratory, can be modified by a number of contextual factors, and have been exceptionally well characterized (Miller and Wang, 2006; Miller et al., 2009, 2010; Roy et al., 2011). The small size of these animals also makes them ideal candidates for freely moving wireless recordings (Miller et al., 2015; Walker et al., 2021; Wong et al., 2023).

Here, we investigated the role of area 32 neurons in vocal communication by implanting chronically an ultra-high density Neuropixels probe (Jun et al., 2017) in area 32 and recording single neurons in freely moving marmosets engaged in bouts of antiphonal calling behaviour. Consistent with the notion that this area acts as an audiovocal interface, we found that a large proportion of neurons exhibited robust modulations of discharge rates in response to externally presented vocalizations as well as before, during, and after self-generated phee calls. We additionally observed varying dynamics of excitation and suppression between presented and produced calls suggestive of an involvement in differentiating perceived and produced vocalizations in social contexts.

## Materials and Methods

### Subjects

Two adult male common marmosets (*Callithrix jacchus*) participated in this study (Marmoset R, age 33 months, weight 463g; Marmoset A, age 49 months, weight 374g). Both animals were under the close supervision of veterinarians throughout their participation. All experiments were performed in compliance with the Canadian Council on Animal Care policy on the care and use of laboratory animals, and the experimental protocol was approved by the Animal Care Committee of the University of Western Ontario Council on Animal Care.

### Surgical Preparation of Animals for Electrophysiological Recordings

In preparation for electrophysiological recordings, each marmoset underwent an aseptic surgical procedure with the dual purpose of creating trephinations in the skull to allow access to the cortex, and fixing a recording chamber to the skull. A microdrill was used to create an approximately 2mm trephination in the right hemisphere above area 32 and lateral to the sagittal sinus, based on subject-specific preoperative anatomical MRI scans and stereotaxic coordinates (Paxinos & Watson & Petrides & Rosa & Tokuno, 2012) The trephination was then sealed with silicone (Kwik-Sil, WPI International, Sarasota, FLA, USA). A second trephination was made roughly 10mm from this location within which a gold amphenol contact (FST inc.19003, Foster City, CA) was secured with dental adhesive for use as an electrical ground in the electrophysiological recordings. A recording chamber was fixed to the skull using a combination of universal dental adhesive (All-Bond Universal, Curion, Richmond, BC, Canada) and UV-cured dental resin cement (Core-Flo DC, Curion, Richmond, BC, Canada), and covered with a protective cap. This chamber provided controlled access to the trephinations, and allowed for stabilization of the head during electrode insertion and implantation. Detailed descriptions of these procedures have been reported previously (Johnston et al., 2018; Schaeffer et al., 2019).

### Vocal Recording and Preparation of Auditory Stimuli

To prepare exemplar calls for use as auditory stimuli for this experiment, we recorded bouts of antiphonal calling between animals in our colony and extracted individual phee calls from these bouts for broadcast to the experimental animals during subsequent combined neural and vocal recording sessions. Within each vocal recording session, one pair of animals was transported from their homeroom to a separate recording room. We used this procedure rather than recording directly in the colony room as the elimination of background noise enabled us to more easily isolate individual calls and attribute them to an identified caller. Animals were placed into separate cages measuring 50.8cm x 60.96cm, 91.44cm, and allowed to move freely. The view of the other cage was obstructed. Vocalizations were recorded by directing a microphone (Sennheiser electronic SE & Co MKH 8050, supercardioid pickup pattern) coupled to a MacBook Pro laptop (2021, running on macOS Ventura 13) via an 18V phantom power (NEEWER model NW100, NEEWER Canada) at one of the two cages. Over the course of each vocal recording session animals spontaneously engaged in multiple bouts of antiphonal calling. We conducted a total of two vocal recording sessions with two pairs of animals. Both pairs were bonded cage-mates. The duration of each session was approximately 120 minutes, within which 675 calls were recorded (32 from pair 1, 643 from pair 2). From each recording session we then extracted individual phee calls using audio editing software (Audacity 3.0, Muse Group). We extracted a total of 34 phee calls (14 from pair 1, 20 from pair 2) and saved these as individual wav files for broadcast to the animals during this experiment.

Prior to electrode implantation and neurophysiological recordings, marmosets R and A were acclimated to the recording room. As our aim was to investigate the role of area 32 in auditory processing and vocal production, we presented the animals with individual phee calls from the aforementioned recordings during this time in order to identify calls which tended to elicit vocal responses from the experimental animals. As a result, a set of 4 phee calls from marmoset S of pair 2, an unfamiliar marmoset to both experimental animals, were selected as stimuli for combined neural and vocal recording sessions.

### Electrophysiological Localization of area 32 and Implantation of Chronic Neuropixels Probes

To allow unrestricted movement during electrophysiological recordings, enabling marmosets to engage in more naturalistic vocal communication, we recorded the activity of single neurons using Neuropixels 1.0 short probes (Jun et al., 2017) implanted chronically within area 32. Prior to implantation, we first identified area 32 based on location and depth from the cortical surface in separate recording sessions in which the animals were restrained as in traditional electrophysiological recording experiments. Detailed descriptions of these procedures have been published previously (Gilliland et al., 2024). These locations were confirmed at the end of the experiments by *ex vivo* ultra high-field structural MRI (see *Confirmation of Recording Sites with ex-vivo MRI*, below)

After localizing area 32, we conducted a single chronic implantation session in each animal, in which we lowered a single Neuropixels 1.0 NHP probe into place at the optimal location determined during the localization sessions, in order to permit freely moving datalogger neural and vocal recordings. These implantation sessions were conducted identically to localization sessions. Neural activity was monitored while advancing the electrode as noted above, and once it had reached a location within area 32 at which we observed well-isolated neurons we allowed it to settle for 30 minutes. Following this, we fixed the electrode in place using a two-step process. We first carefully flowed silicone (Kwik-Sil, WPI International, Sarasota, FLA, USA) around the electrode shank within the trephination until it was completely sealed and the shank was fully encased. We then covered this with UV-cured dental resin cement (Core-Flo DC, Curion, Richmond, BC, Canada) until the probe was secured up to roughly 5mm above the skull surface. Once the cement was cured, we carefully detached the custom electrode holder from the probe and slowly raised it using the stereotaxic micromanipulator. Once this was done, we added additional layers of cement to fully encase the probe base. Following this, we further cemented in place around the electrode a custom-designed, 3D printed protective cone/headstage holder. This served the dual purpose of protecting the site of electrode implantation as well as providing an anchor point for the Spike Gadgets headstage and datalogger (Spike Gadgets, SanFrancisco, CA).

### Simultaneous Vocal and Neural Recordings in Freely Moving Marmosets During Phee Call Broadcasts

For each recording session, the animal was transported to the recording room and prepared for untethered datalogger recordings. To do this, we attached the SpikeGadgets Neuropixels datalogger headstage to the previously implanted headstage holder and commenced recording. Neural data were recorded on a microSD card inserted into the headstage (Samsung PRO Plus Micro SD 256GB). The Spike Gadgets system was configured to receive both analog inputs from a small lavalier microphone used to record vocalizations (Hollyland Technology Lark M1, Shenzhen, Guandong, China) and digital sync pulses from a Raspberry Pi 3 Model B, which was used to broadcast the previously recorded calls via a Bose Soundlink III speaker (Bose Corp, Framingham Mass.) placed a distance of 1m from the animal. These inputs were directed to the SpikeGadgets environmental control system (ECU), which served as an interface to the SpikeGadgets MCU, and enabled synchronization of played and recorded calls with neural data. Neural recordings were controlled by the SpikeGadgets Trodes software package. After attaching the headstage and initiating wireless recording, the animal was released into a 30cm x 20cm x 30cm transfer box with clear sides allowing full visibility of the recording lab, and within which they were allowed unrestricted movement. The lavalier microphone was positioned at a small opening at the top of the box to sample vocalizations. Auditory stimuli consisting of the previously recorded phee calls were played manually under the control of the experimenter, triggered by keystrokes via custom written Python code on the Raspberry pi. Dynamics of stimulus presentation were intended to mimic that of natural antiphonal calling. In some cases call sequences were initiated by calls broadcast by the experimenter, while in others spontaneous calls evoked by the animals were “answered” by the experimenter. Response calls were initiated within 6 seconds of calls produced by the animal, based on natural antiphonal calling dynamics (Miller and Wang, 2006). Each session lasted approximately 30-45min. The duration of each session was dictated by the animal’s willingness to respond to played calls or generate spontaneous calls, and was ended by the experimenter.

### Automated Spike Sorting with Manual Curation

Spike sorting was performed using Kilosort4 (Pachitariu et al., 2016) followed by manual curation with Phy. Preprocessing involved median filtering and whitening, followed by adaptive template matching. Single-unit classification was based on waveform consistency, amplitude distribution, autocorrelogram characteristics, and unit stability. Detailed descriptions of this process have been reported previously (Gilliland et al., 2024; Selvanayagam et al., 2024).

### Automated Detection of Phee Calls

Times of call onset and offset were determined using a band-limited entropy detector in the Raven Pro software program (Cornell Lab of Ornithology, Ithaca NY, https://www.ravensoundsoftware.com/software/raven-pro/) and were additionally visually inspected and manually corrected as needed.

### Confirmation of Recording Sites With Ex-vivo MRI

Following data collection, the marmosets were perfused, and electrode tracts were reconstructed using ex-vivo ultra-high field MRI. Animals were deeply anesthetized, perfused transcardially, and heads were post-fixed in 10% buffered formalin. MRI scans were conducted at 15.2T (Bruker BioSpec Avance Neo) using a 35-mm quadrature-detection volume coil. T2*-weighted images (75 × 75 × 75 µm resolution) were acquired and registered to the NIH ex-vivo marmoset brain atlas (Liu et al., 2018). These procedures have been described in detail previously (Gilliland et al., 2024) ***Experimental Design and Statistical Analysis***

All analyses were performed using scripts custom-written in Matlab (Mathworks, Natick MA, USA). The analysis pipeline was designed to extract, process, and interpret neural data aligned to both the presentation of phee calls and the onset of produced phee calls. The first vocalization occurring after each presented phee call was identified. Matching trials, along with the time differences between stimulus and vocalization onsets, were stored for further analysis. Neural spike data were loaded from Kilosort-generated files. Spike times were converted to milliseconds and aligned to two reference points: presented phee call onset and produced vocalization onset. For each neuron, a binary spike matrix was generated, representing spike occurrence across trials and time.

To assess neuronal responsiveness, spike counts in three phases—stimulus (0-1000 ms after phee call presentation), pre-vocalization (-2000 to 0 ms prior to produced vocalization), and post-vocalization (0-1000 ms after produced vocalization)—were compared to baseline activity using Wilcoxon signed-rank tests at p<0.05. Neurons were classified based on significant activity changes in one or more phases. Categories included stimulus-only, pre-vocalization-only, post-vocalization-only, combinations of these phases, and non-responsive neurons. The proportions of neurons in each category were calculated and visualized in pie charts for each of the two sessions, providing an overview of the neural population’s functional diversity. Spike counts were aggregated and smoothed using a 100 ms Gaussian kernel to generate PSTHs to illustrate the temporal firing patterns of neurons relative to call presentation and call production. Principal component analysis (PCA) was applied to z-normalized neural activity aligned to call presentation (-1000 ms to 1500ms) and call production (-1000 to 1500 ms) to reduce data dimensionality and identify dominant patterns of activity. The first two PCA from the activity aligned to call presentation and the PCA from the activity aligned on call production were used as input features for k-means clustering which was run 1000 times. The optimal number of clusters was identified using silhouette analysis, which evaluates the consistency and separation of clusters. Neurons in each cluster were visualized using scatter plots of PC1 with the activity aligned to call presentation and PC1 with the activity aligned on call production. Average z-normalized neuronal waveforms for each cluster, aligned to both call presentation and call production, were plotted with standard error of the mean. To visualize the activity pattern of all single neurons in the identified clusters, we generated heatmaps of z-normalized neural activity binned into 25 ms internals and aligned on call presentation and call production. Neurons were sorted by the time to reach 50% of their maximal activity for call presentation or call production. To investigate neural response differences between presented and generated vocalizations, we performed a classification analysis using raw spiking data in 1s windows immediately after the presentation of calls and in the 1s after the production of calls. Data were split into training (50%) and testing (50%) sets. A support vector machine (SVM) classifier with a linear kernel was trained on the training set and tested on the held-out data. This process was repeated 1000 times to assess classification accuracy under varying splits of the data. Classification accuracy was calculated for each iteration as the proportion of correctly predicted labels in the test set. A violin plot was used to display the distribution of classification accuracies across iterations.

## Results

We recorded neural activity from the ACC of two freely moving marmoset monkeys using high-density Neuropixels probes implanted in each animal coupled with a SpikeGadgets datalogger. During these sessions, we presented each marmoset with previously recorded conspecific phee calls while also tracking their self-generated phee calls (Fig. 1A). To ensure precise identification of vocalization times, we utilized the Raven Pro 1.6 software developed by the Cornell Laboratory of Ornithology, which allowed us to automatically detect and subsequently confirm by visual inspection the onset and offset times of phee calls produced by the animals from which neural recordings were obtained (Fig. 1B). Marmoset R typically responded with a phee call approximately 2 seconds after hearing the presented call, displaying a relatively consistent response time. In contrast, marmoset A exhibited much greater variability in call onset times following the presented vocalizations, indicating individual differences in response patterns (Fig. 1C). The yield of recorded neurons from the implanted Neuropixels probes differed between the two animals (Fig. 1D). Although both animals initially exhibited a similar number of isolated neurons—125 in marmoset R and 129 in marmoset A, recorded a few hours post-implantation— the neuron count declined in marmoset R to 17 on day 1 and 16 on day 2. In contrast, the neuron count in marmoset A increased to over 200 for the subsequent three days. To prevent re-analysis of the same neurons across sessions, we included only one session from marmoset R (day 0, n = 125 neurons) and marmoset A (day 1, n = 204 neurons) in our final analysis, verifying that results remained consistent across all sessions for marmoset R. Ex-vivo ultra-high field magnetic resonance imaging at 15.2 Tesla confirmed that the probes primarily targeted areas 32 and 32v of the anterior cingulate cortex, with a few neurons recorded in area 14R in marmoset A (Figs. 1E, F)

**Figure 1.**
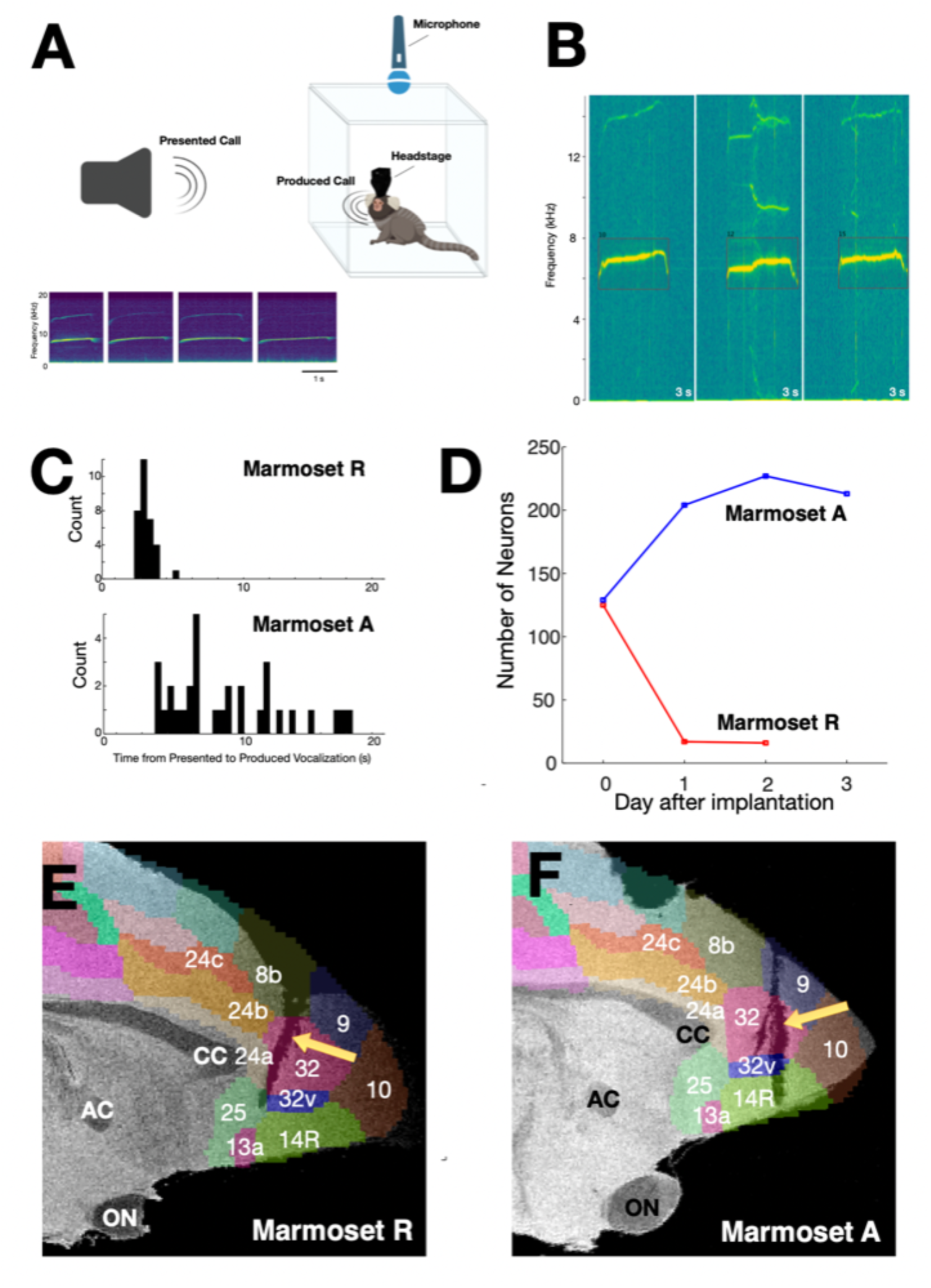
Experimental setup, vocal response behavior, neuronal yield, and implant location in marmosets. **(A)** Schematic of the experimental setup. Neural recordings from freely moving marmosets were captured using high-density Neuropixels probes implanted in the anterior cingulate cortex, with data logged via a SpikeGadgets datalogger. The marmosets were presented with conspecific phee calls while their own vocalizations were also tracked. **(B)** Illustration of precise detection of vocalization onsets and offsets using Raven Pro. **(C)** Histogram of response times from presented to produced phee calls in marmosets R and A. **(D)** Number of neurons recorded over three days post-implantation for each marmoset. **(E, F)** Ex-vivo MRI scans at 15.2T showing electrode tracts (yellow arrows) in the anterior cingulate cortex for marmosets R (E) and A (F). The overlaid color-shaded brain areas (Paxinos et al., 2012) illustrate targeted and adjacent cytoarchitectonic areas. AC, anterior commissure; CC, corpus callosum; ON, optic nerve.

We observed a variety of neural response profiles in the ACC. Consistent with our previous findings that many area 32 neurons in marmosets respond to conspecific vocalizations, we found neurons in this naturalistic vocalization paradigm that responded to the presented phee calls. Figure 2A illustrates a neuron that exhibited a transient increase in activity following the presented phee calls (blue triangles), which returned to baseline levels before the marmoset produced its own phee calls (magenta circles). A similar pattern is seen in Figure 2B, where the neuron also shows some activity following the onset of the produced calls (magenta circle, right panel).

**Figure 2.**
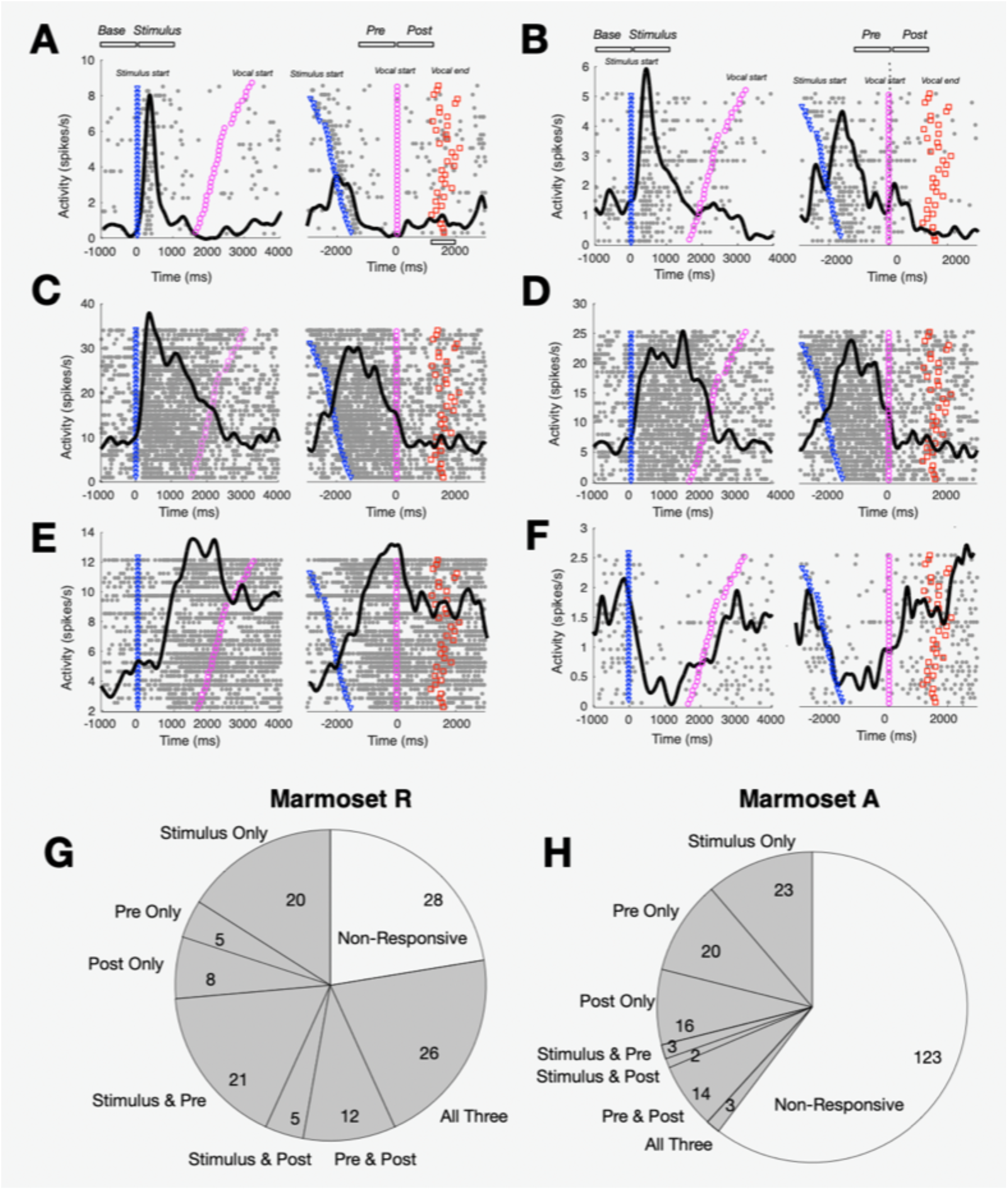
Neural activity patterns of significantly modulated neurons in response to presented and produced vocalizations in marmosets R and A. **(A-F)** Raster plots and peristimulus time histograms (PSTHs) showing activity of single neurons aligned to presented (stimulus) and self-generated (vocalization) phee calls. Each panel displays activity during four time periods: baseline (Base), stimulus presentation (Stimulus), pre-vocalization (Pre), and post-vocalization (Post). Rasters depict spike times (gray dots) across trials, with color-coded markers indicating specific event timings: stimulus start (magenta circles), vocal start (blue triangles), and vocal end (red squares). Black lines indicate the average spike density across trials. **(G, H)** Proportion of neurons with significant responses to presented calls, produced calls, or both, in marmoset R (G) and marmoset A (H). Pie charts categorize neurons as responsive to stimulus only, pre-vocalization only, post-vocalization only, combinations of these periods (Stimulus & Pre, Stimulus & Post, Pre & Post), all three periods, or non-responsive.

Some neurons displayed more sustained responses to presented phee calls, only decreasing just before the onset of the produced vocalization (Fig. 2C), and these neurons did not show increased activity during the produced calls above baseline levels. Other neurons, such as the one shown in Figure 2D, increased their activity following the presented phee calls but reached peak activity at a later time. Additionally, many neurons showed increased activity just before and during the produced phee calls, peaking at or after call onset (Fig. 2E). In contrast, some neurons reduced their activity following the presented call and showed an increase immediately prior to the produced call (Fig. 2F).

To quantify the total number of neurons modulated during this free vocalization paradigm, we identified neurons with significant activity changes across different task epochs, compared to baseline activity evaluated in a window from 1000-0 ms before phee call presentation. We used nonparametric Wilcoxon signed-rank tests (P < 0.05) to test for significance. Modulated neurons were found in all three task periods analyzed: the stimulus period (0-1000 ms after phee call presentation), the pre-vocalization period (2000-0 ms before the produced call), and the post-vocalization period (0-1000 ms after the produced call) (Figs. 1G and H). Note that the stimulus and pre-vocal periods overlapped somewhat for neurons recorded from marmoset R, which typically responded with a call 2-3 seconds after phee call presentation. Overall, 78% of neurons in marmoset R and 40% of neurons in marmoset A showed significant modulation across these broad task periods.

To examine the neural response dynamics during both presented and self-generated vocalizations, we performed principal component analysis (PCA) on the 178 task-modulated neurons from the two monkeys. Figure 3 illustrates the temporal evolution of the top three principal components (PC1, PC2, and PC3) during presented (Fig. 3, left panels) and produced (Fig. 3, right panels) phee calls, capturing key aspects of neural response variance in each condition.

**Figure 3.**
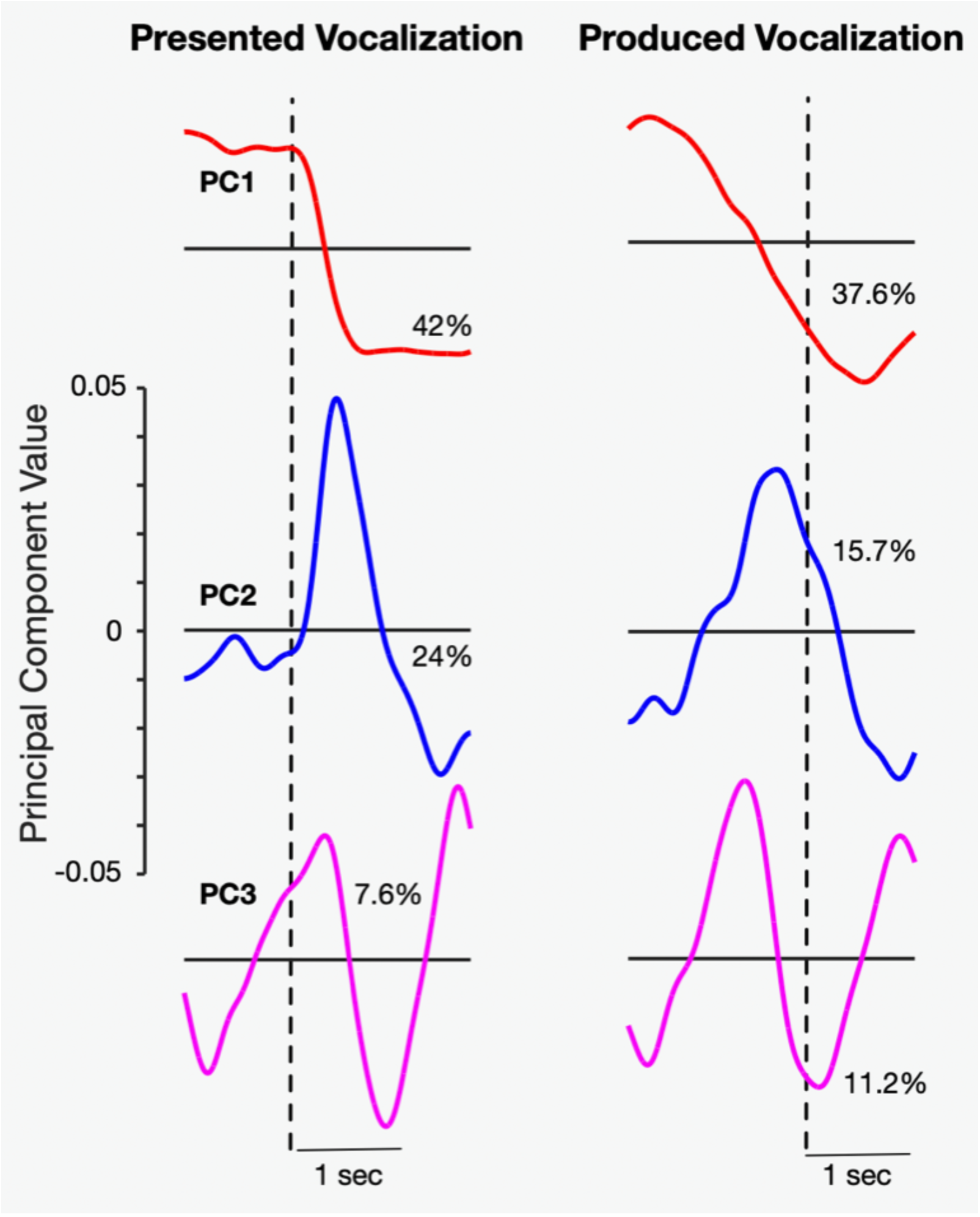
Principal component analysis of neural responses to presented and produced vocalizations. (**Left Panels)** Principal component (PC) analysis of neural activity during presented vocalizations. The time courses of the first three principal components (PC1, PC2, and PC3) are shown. The percentage next to each component indicates the variance explained by that component. **(Right panels)** Principal component analysis of neural activity during produced vocalizations.

In response to the presented phee calls (Fig. 3, left panels), the first principal component (PC1) showed a marked decrease in activity immediately following the onset of the call, with a sustained level afterwards. This component explained 42% of the variance in the neural response, suggesting it captures a significant portion of the activity in area 32 associated with external vocal stimuli. The second principal component (PC2), which accounted for 24% of the variance, exhibited a phasic increase in activity following the call onset. The third principal component (PC3) displayed a more complex sequence of modulation, and explained 7.6% of the variance.

During self-generated phee calls (Fig. 3, right panels), neural dynamics differed noticeably. PC1 showed a decrease in activity immediately after the onset of the produced call, capturing 37.6% of the variance. This response pattern was consistent with the decrease observed in response to presented calls. PC2, which explained 15.7% of the variance, exhibited a transient increase starting before call onset, consistent with activation during vocal production. Similar to its response profile during presented calls, PC3 displayed an oscillatory pattern, explaining 11.2% of the variance, suggesting this component may capture a more general response pattern present across both presented and produced vocalization conditions.

To classify distinct neural response patterns during presented and produced vocalizations, we used k-means clustering analysis on the first principal component of the activity for presented vocalizations and produced vocalizations of the 178 significantly modulated neurons. Using silhouette analysis to determine the optimal number of clusters, we observed a peak average silhouette value at 3 clusters (Fig. 4 A), indicating that the modulated neurons can be grouped into three distinct response types based on their activity patterns during presented and produced vocalizations.

**Figure 4.**
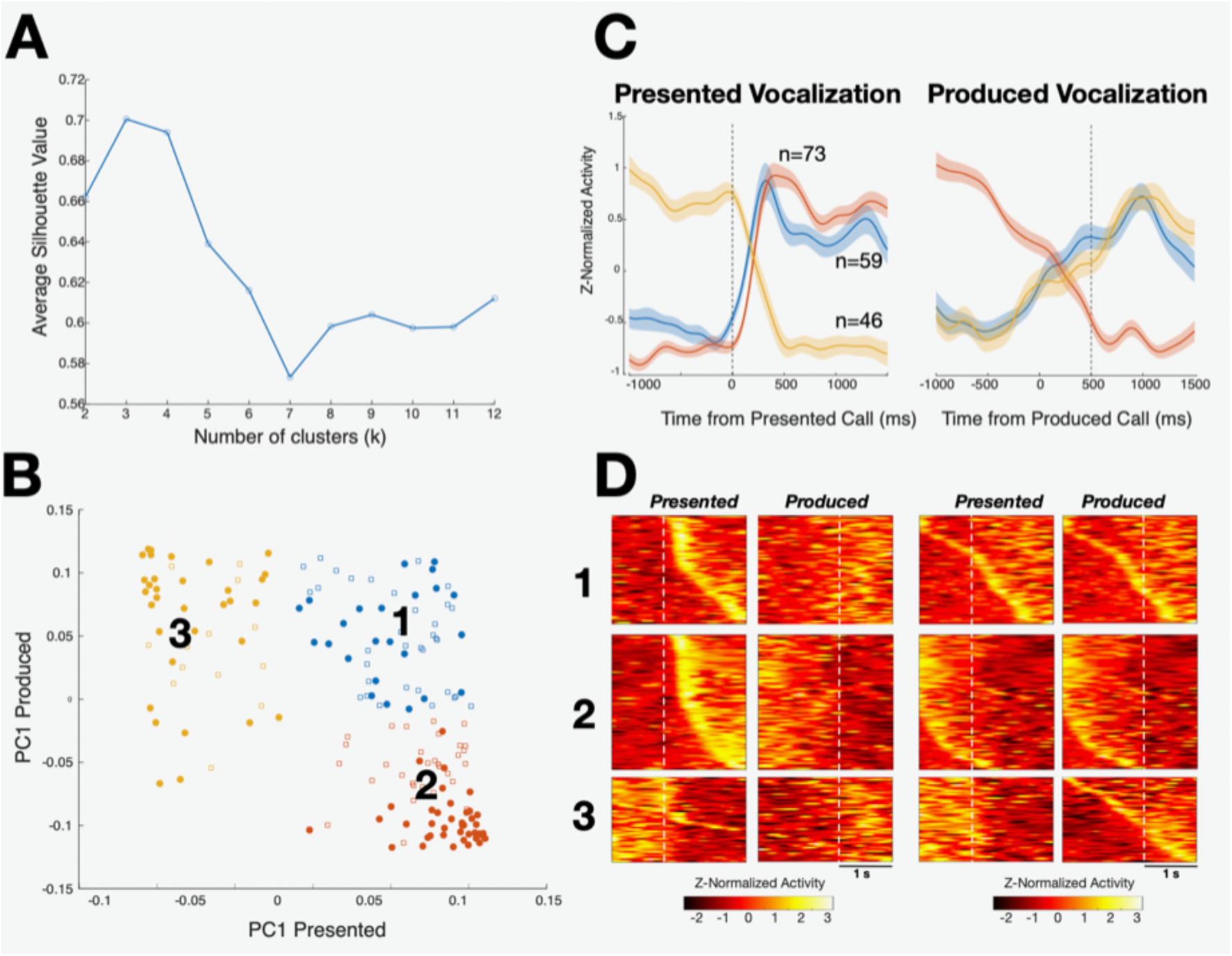
Clustering analysis of neural response patterns to presented and produced vocalizations. **(A)** Silhouette analysis showing the average silhouette value as a function of the number of clusters (k) for k-means clustering. A peak at 3 clusters indicates that the neural activity can be optimally divided into three distinct response types. **(B)** Scatter plot of principal component 1 (PC1) scores for presented versus produced vocalizations, with each point representing a neuron. Colors denote the three identified clusters. Each cluster contains neurons from both marmosets R (filled circles) and A (open circles). **(C)** Average z-normalized neural activity of each identified cluster during presented (left) and produced (right) phee calls. Shaded regions indicate ±SEM. The dotted line marks the onset of the presented call (left) and the produced call (right). **(D)** Heat maps showing the z-normalized neuronal activity for neurons in the 3 clusters during call presentation and call production. (*Left*) Each heat map represents neuronal activity sorted by the time at which activity reaches 50% of the maximal response for call presentation. Data are aligned to the onset of the presented and produced calls (vertical dashed lines at 0 ms). Rows represent individual neurons within each cluster. The consistent sorting across presented and produced calls reveals distinct activity patterns in different clusters. (*Right*) Same as presented at left, but sorted by the time at which activity reached 50% of the maximal response for call production.

In Figure 4B, we plot each neuron’s principal component 1 scores for presented versus produced vocalizations. The scatter plot shows the three identified clusters, with each cluster containing neurons from both marmosets (R and A, open and filled circles, respectively), represented by different colors. The separation of clusters for presented and produced vocalizations suggests that distinct neural populations are selectively tuned to either vocalization source or exhibit different response magnitudes and directions based on the vocalization type.

Figure 4C shows the average z-normalized neural activity for each of the three clusters during presented (left) and produced (right) phee calls. Figure 4D shows the z-normalized neural activity of single neurons in the clusters. The dotted lines indicate the onset of the presented phee call (left panels) and the onset of the produced phee call (right panels). Neurons in cluster 1 (red, n=73) exhibited a marked sustained increase in activity following the onset of the presented phee call and decreased their activity before the onset of the produced call. This pattern suggests that these neurons respond strongly to external vocalizations but are suppressed during self-generated vocalizations, indicating a possible role in distinguishing between external auditory inputs and self-generated actions. Cluster 2 (blue, n=59) showed a phasic increase in activity after the presented call, followed by a gradual increase that began approxinately 500 ms before the onset of the produced call and peaked about 1000 ms afterward. This response pattern implies that these neurons are inhibited by external vocalizations but become active in preparation for and during self-generated calls, indicating a potential role in vocal initiation. Cluster 3 (yellow, n=46) displayed a decrease in activity after the presented calls but like cluster 3 increased its activity before and during produced vocalizations.

To further evaluate the ability to distinguish between neural activity patterns associated with presented versus produced calls, we trained a linear Support Vector Machine (SVM) classifier using the activity of all recorded neurons within each of the two sessions. For each trial, neural responses were represented by the summed spike counts of each neuron within the first second following the onset of either a presented or generated call, creating a high-dimensional feature space that captured the collective neural response dynamics for each vocalization type. This analysis aimed to quantify the separability of neural responses on a trial-by-trial basis, leveraging the simultaneous activity patterns across the neural population to enhance classification accuracy.

The classifier demonstrated a mean correct classification rate of 96% for marmoset R, indicating strong separability between the neural responses associated with presented and generated calls, with a distinct pattern of activity following each call type. For marmoset A, the classifier achieved a mean correct classification rate of 82%, suggesting effective, albeit slightly lower, discriminability in neural response patterns compared to marmoset R. These findings highlight the high degree of specificity in neural encoding for presented versus self-generated calls within each marmoset’s neural population in area 32, indicating robust neural differentiation based on the source of the vocalization.

## Discussion

Convergent lines of anatomical and functional evidence have linked area 32 with both auditory processing and vocal production. The confluence of these processes within this area suggest that it may act as an audiovocal interface linking auditory inputs to vocal outputs. Given the well-established links between pregenual ACC and social cognition (see for review Amodio and Frith, 2006), we reasoned that this may be particularly the case in contexts requiring voluntary control of vocal initiation on the basis of cognitive factors, such as during communication with a conspecific (Jürgens, 2009; Fichtel and Manser, 2010; Bradbury and Vehrencamp, 2011; Grijseels et al., 2023). Here, we investigated this by carrying out electrophysiological recordings in freely moving marmosets engaged in an antiphonal calling paradigm in which they responded to broadcasts of long-distance phee calls by voluntarily generating phee calls (Miller and Wang, 2006; Eliades and Miller, 2017). We found that the response properties of single neurons in marmoset area 32 did indeed meet the basic criteria for an area integrating auditory information with vocal output, and potentially cognitive factors. First, consistent with our prior finding of complex response dynamics in the auditory responses of area 32 neurons to biological sounds and conspecific vocalizations (Gilliland et al., 2024) a large proportion of neurons we recorded here (58.9 % of responsive neurons across both animals) were modulated by presented phee calls in the context of the antiphonal calling paradigm. Second, roughly 80% of responsive neurons were modulated around the time at which marmosets generated calls, consistent with prior findings suggestive of role in call production (Jürgens and Ploog, 1970; Sutton et al., 1974). Third, nearly all responsive neurons exhibited either a sustained increase or suppression of activity in the interval between presented and produced vocalizations. Such modulations, which bridge the temporal gap between sensory responses and motor output have been associated previously with varied cognitive processes including working memory (Pasternak and Greenlee, 2005) and decision making (Shadlen and Kiani, 2013), as well as motor preparation (Jonikaitis et al., 2023). It must be noted however that since we did not here explicitly manipulate any cognitive or motor preparatory processes, such a link must remain speculative (Teller, 1984) and awaits further investigation.

Our findings of robust responses of area 32 neurons to presented and produced vocalizations are consistent with prior conceptualizations of the role of the ACC in vocal control. On the basis of stimulation and lesion studies targeting ACC and the interconnected PAG (Müller-Preuss and Jürgens, 1976), a structure with direct connections to the brainstem vocal pattern generator (Mantyh, 1983), Jurgens (2009) proposed that the ACC, including an area corresponding to area 32, existed at the apex of a hierarchical cingulo-periaqueductal pathway involved in voluntary control of vocalizations. In a similar vein, the dual-network model of Hage & Neider (2016) places the ACC, including area 32, as the highest level of a limbic vocal-initiating network which includes the PAG, drives the vocal pattern generator in the brainstem, and additionally receives top-down inputs from a suite of lateral prefrontal, premotor, and motor cortical areas responsible for high-level vocal control. In this conceptualization the ACC is in a sense a linchpin connecting evolutionarily old structures critical for emotional vocalizations with relatively newer cortical areas involved in cognitive control. Here, we found that area 32 neurons exhibited neural responses during vocal communication that resembled those observed in other areas with respect to vocal production but differed with respect to auditory responsiveness to conspecific calls. Numerous studies in both macaque monkeys performing trained vocalization tasks (Hage and Nieder, 2013, 2016; Gavrilov et al., 2017; Hage, 2018), and marmosets engaged in naturalistic antiphonal calling (Miller et al., 2015; Roy et al., 2016; Nummela et al., 2017; Jovanovic et al., 2022; Zhao and Wang, 2023), have demonstrated modulations in discharge rates of neurons in prefrontal and premotor cortex in advance of and during the production of vocalizations. The complex excitatory and inhibitory discharge dynamics around the time of phee call production during antiphonal calling in marmosets (Miller et al., 2015; Roy et al., 2016), resemble closely those we observed here. In contrast, auditory responses to presented calls in the antiphonal calling paradigm appear to be relatively weak and present in only roughly 10% of neurons in cytoarchitectonic areas corresponding to marmoset premotor cortex (Miller et al., 2015; Roy et al., 2016; Nummela et al., 2017), though there is some evidence that auditory responsiveness may be greater in more anterior frontal cyotoarchitectonic subfields including areas 8aD, 8aV, 46, 47, 9 and 10 (Nummela et al., 2017; Jovanovic et al., 2022; Wong et al., 2024). This differs considerably from the robust auditory responses we observed in area 32 here and in our previous work (Gilliland et al., 2024), suggesting that this area is related more directly to sensory processing of auditory inputs in the context of vocal communication. In the PAG, which as noted above is related more directly to vocal production, auditory responses are almost completely absent (Düsterhöft et al., 2004). Single neurons exhibit exclusively excitatory dynamics at various times prior to and during vocalizations which may be involved in coordinating various motor processes related to vocal production (Larson and Kistler, 1984, 1986; Larson, 1991). Overall, considered in the context of networks responsible for vocal production, our findings are broadly consistent with the conceptualization of area 32 as an “auditory field” (Medalla and Barbas, 2014). This relative bias toward sensory input observed together with activity changes occurring preceding and following vocalization onset suggest additionally that area 32 may implement an early stage of transformation from auditory inputs which are converted to motor or bias signals shared with other cortical and subcortical regions with more direct access to the vocal pattern generator in the brainstem. Indeed, it has been shown that the PAG, as well cortical motor, premotor, and cingulate motor areas, but not area 32, have disynaptic connections to laryngeal motorneurons involved in vocal control (Cerkevich et al., 2022).

As noted above, a large body of evidence from a variety of sources has implicated pregenual ACC, including area 32 in a range of emotional and social cognitive functions involving evaluations of the self and others (Amodio and Frith, 2006). In human participants these include self-reflective thoughts (Johnson et al., 2002), monitoring of somatic states (Blakemore et al., 2000; Shergill et al., 2013) and evaluations of the self with respect to unknown others (Ochsner et al., 2005). In nonhuman primates, the functional connectivity of this region has been shown to be related to social group size (Sallet et al., 2011), and it has been linked to the valuation of social information (Rudebeck et al., 2006), encoding of social behaviors (Van Mao et al., 2017), and the regulation of decision making on the basis of emotional states (Amemori and Graybiel, 2012; Wallis et al., 2017, 2019). In the context of antiphonal calling in the marmoset model, the dynamics of vocal communication are affected profoundly by emotional and social factors as arousal state (Borjon et al., 2016), call timing (Miller and Wang, 2006; Miller et al., 2009), social context (Jovanovic et al., 2022) and caller identity (Miller and Wren Thomas, 2012), and one recent model of antiphonal calling behavior has incorporated many of these factors in the prediction of vocal turn-taking behaviour (Grijseels et al., 2024). Given the demonstrated encoding of context in lateral prefrontal areas during antiphonal calling (Nummela et al., 2017; Jovanovic et al., 2022), the established link between pregenual ACC and social and cognitive processes, it seems reasonable to suggest that the modulations in activity we observed in area 32 may additionally reflect aspects of social cognition related to the probability and timing of call production. Here our current data are equivocal as in this study our primary aim was to characterize basic response properties in area 32 and we therefore did not systematically manipulate such factors. Future studies incorporating social cognitive factors in this model system have the promise to reveal whether and how area 32 integrates social information and emotional states with incoming auditory signals to influence call production during naturalistic vocal communication.

## Acknowledgments

We wish to thank Cheryl Vander Tuin, Whitney Froese, and Hannah Pettypiece for animal preparation and care and Joseph Gati for scanning assistance. Support was provided by the Canadian Institutes of Health Research (S.E.), and the Natural Sciences and Engineering Council of Canada (S.E.). We also acknowledge the support of the Government of Canada’s New Frontiers in Research Fund (NFRF) [NFRF-T-2022-00051].

## References

Amemori KI, Graybiel AM (2012) Localized microstimulation of primate pregenual cingulate cortex induces negative decision-making. Nat Neurosci 15:776–785 Available at: https://pubmed.ncbi.nlm.nih.gov/22484571/ [Accessed February 11, 2025].

Amodio DM, Frith CD (2006) Meeting of minds: the medial frontal cortex and social cognition. Nat Rev Neurosci 7:268–277 Available at: https://pubmed.ncbi.nlm.nih.gov/16552413/ [Accessed February 11, 2025].

Bezerra BM, Souto A (2008) Structure and usage of the vocal repertoire of Callithrix jacchus. Int J Primatol 29:671–701 Available at: https://link.springer.com/article/10.1007/s10764-008-9250-0 [Accessed February 11, 2025].

Blakemore SJ, Smith J, Steel R, Johnstone EC, Frith CD (2000) The perception of self-produced sensory stimuli in patients with auditory hallucinations and passivity experiences: evidence for a breakdown in self-monitoring. Psychol Med 30:1131–1139 Available at: https://pubmed.ncbi.nlm.nih.gov/12027049/ [Accessed February 11, 2025].

Borjon JI, Takahashi DY, Cervantes DC, Ghazanfar AA (2016) Arousal dynamics drive vocal production in marmoset monkeys. J Neurophysiol 116:753–764 Available at: https://pubmed.ncbi.nlm.nih.gov/27250909/ [Accessed February 11, 2025].

Bradbury JW, Vehrencamp SL (2000) Economic models of animal communication. Anim Behav 59:259–268 Available at: https://pubmed.ncbi.nlm.nih.gov/10675247/ [Accessed February 11, 2025].

Bradbury JW, Vehrencamp SL (2011) Principles of animal communication, 2nd ed.: 697 Available at: https://global.oup.com/academic/product/principles-of-animal-communication-9780878930456 [Accessed February 11, 2025].

Cerkevich CM, Rathelot JA, Strick PL (2022) Cortical basis for skilled vocalization. Proc Natl Acad Sci U S A 119:e2122345119 Available at: https://www.pnas.org/doi/abs/10.1073/pnas.2122345119 [Accessed January 10, 2023].

Dureux A, Zanini A, Everling S (2024) Mapping of facial and vocal processing in common marmosets with ultra-high field fMRI. Commun Biol 7 Available at: https://pubmed.ncbi.nlm.nih.gov/38480875/ [Accessed October 1, 2024].

Düsterhöft F, Häusler U, Jürgens U (2004) Neuronal activity in the periaqueductal gray and bordering structures during vocal communication in the squirrel monkey. Neuroscience 123:53–60.

Eliades SJ, Miller CT (2017) Marmoset vocal communication: Behavior and neurobiology. Dev Neurobiol 77:286–299 Available at: https://pubmed.ncbi.nlm.nih.gov/27739195/ [Accessed July 23, 2023].

Fichtel C, Manser M (2010) Vocal communication in social groups. Animal Behaviour: Evolution and Mechanisms:29–54 Available at: https://link.springer.com/chapter/10.1007/978-3-642-02624-9_2 [Accessed February 11, 2025].

Gavrilov N, Hage SR, Nieder A (2017) Functional Specialization of the Primate Frontal Lobe during Cognitive Control of Vocalizations. Cell Rep 21:2393–2406 Available at: https://pubmed.ncbi.nlm.nih.gov/29186679/ [Accessed March 22, 2023].

Gilliland RL, Selvanayagam J, Zanini A, Johnston KD, Everling S (2024) Neural activity for complex sounds in the marmoset anterior cingulate cortex. Commun Biol 7:1310 Available at: https://pubmed.ncbi.nlm.nih.gov/39394433/ [Accessed January 23, 2025].

Grijseels DM, Prendergast BJ, Gorman JC, Miller CT (2023) The neurobiology of vocal communication in marmosets. Ann N Y Acad Sci 1528 Available at: https://pubmed.ncbi.nlm.nih.gov/37615212/ [Accessed February 5, 2024].

Hadland KA, Rushworth MFS, Gaffan D, Passingham RE (2003) The effect of cingulate lesions on social behaviour and emotion. Neuropsychologia 41:919–931 Available at: https://pubmed.ncbi.nlm.nih.gov/12667528/ [Accessed February 11, 2025].

Hage SR (2018) Auditory and audio-vocal responses of single neurons in the monkey ventral premotor cortex. Hear Res 366:82–89 Available at: https://pubmed.ncbi.nlm.nih.gov/29598839/ [Accessed August 7, 2022].

Hage SR, Nieder A (2013) Single neurons in monkey prefrontal cortex encode volitional initiation of vocalizations. Nat Commun 4 Available at: https://pubmed.ncbi.nlm.nih.gov/24008252/ [Accessed February 11, 2025].

Hage SR, Nieder A (2016) Dual Neural Network Model for the Evolution of Speech and Language. Trends Neurosci 39:813–829 Available at: https://pubmed.ncbi.nlm.nih.gov/27884462/ [Accessed March 22, 2023].

Ito S, Stuphorn V, Brown JW, Schall JD (2003) Performance monitoring by the anterior cingulate cortex during saccade countermanding. Science 302:120–122 Available at: https://pubmed.ncbi.nlm.nih.gov/14526085/ [Accessed February 11, 2025].

Jafari A, Dureux A, Zanini A, Menon RS, Gilbert KM, Everling S (2023) A vocalization-processing network in marmosets. Cell Rep 42:112526 Available at: https://pubmed.ncbi.nlm.nih.gov/37195863/ [Accessed July 2, 2023].

Johnson SC, Baxter LC, Wilder LS, Pipe JG, Heiserman JE, Prigatano GP (2002) Neural correlates of self-reflection. Brain 125:1808–1814 Available at: https://pubmed.ncbi.nlm.nih.gov/12135971/ [Accessed February 11, 2025].

Johnston KD, Barker K, Schaeffer L, Schaeffer D, Everling S (2018) Methods for chair restraint and training of the common marmoset on oculomotor tasks. J Neurophysiol 119:1636–1646.

Jonikaitis D, Noudoost B, Moore T (2023) Dissociating the Contributions of Frontal Eye Field Activity to Spatial Working Memory and Motor Preparation. J Neurosci 43 Available at: https://pubmed.ncbi.nlm.nih.gov/37871965/ [Accessed February 11, 2025].

Jovanovic V, Fishbein AR, de la Mothe L, Lee KF, Miller CT (2022) Behavioral context affects social signal representations within single primate prefrontal cortex neurons. Neuron 110:1318–1326.e4 Available at: https://pubmed.ncbi.nlm.nih.gov/35108498/ [Accessed August 6, 2022].

Jun JJ et al. (2017) Fully integrated silicon probes for high-density recording of neural activity. Nature 551:232–236 Available at: https://pubmed.ncbi.nlm.nih.gov/29120427/ [Accessed January 24, 2022].

Jürgens U (1994) The role of the periaqueductal grey in vocal behaviour. Behavioural brain research 62:107–117 Available at: https://pubmed.ncbi.nlm.nih.gov/7945960/ [Accessed February 11, 2025].

Jürgens U (2009) The Neural Control of Vocalization in Mammals: A Review. Journal of Voice 23:1–10.

Jürgens U, Ploog D (1970) Cerebral representation of vocalization in the squirrel monkey. Exp Brain Res 10:532–554 Available at: https://pubmed.ncbi.nlm.nih.gov/4988409/ [Accessed February 11, 2025].

Kolling N, Behrens TEJ, Wittmann MK, Rushworth MFS (2016) Multiple signals in anterior cingulate cortex. Curr Opin Neurobiol 37:36–43 Available at: https://pubmed.ncbi.nlm.nih.gov/26774693/ [Accessed February 11, 2025].

Larson CR (1991) On the relation of PAG neurons to laryngeal and respiratory muscles during vocalization in the monkey. Brain Res 552:77–86.

Larson CR, Kistler MK (1984) Periaqueductal gray neuronal activity associated with laryngeal EMG and vocalization in the awake monkey. Neurosci Lett 46:261–266 Available at: https://pubmed.ncbi.nlm.nih.gov/6738919/ [Accessed February 11, 2025].

Larson CR, Kistler MK (1986) The relationship of periaqueductal gray neurons to vocalization and laryngeal EMG in the behaving monkey. Exp Brain Res 63:596–606 Available at: https://pubmed.ncbi.nlm.nih.gov/3758271/ [Accessed March 28, 2023].

Liu C, Ye FQ, Yen CCC, Newman JD, Glen D, Leopold DA, Silva AC (2018) A digital 3D atlas of the marmoset brain based on multi-modal MRI. Neuroimage.

Mantyh PW (1983) Connections of midbrain periaqueductal gray in the monkey. II. Descending efferent projections. J Neurophysiol 49:582–594 Available at: https://pubmed.ncbi.nlm.nih.gov/6300351/ [Accessed February 11, 2025].

Medalla M, Barbas H (2014) Specialized prefrontal “auditory fields”: organization of primate prefrontal-temporal pathways. Front Neurosci 8 Available at: https://pubmed.ncbi.nlm.nih.gov/24795553/ [Accessed July 20, 2023].

Medalla M, Lera P, Feinberg M, Barbas H (2007) Specificity in inhibitory systems associated with prefrontal pathways to temporal cortex in primates. Cereb Cortex 17 Suppl 1 Available at: https://pubmed.ncbi.nlm.nih.gov/17725996/ [Accessed February 14, 2023].

Miller CT, Beck K, Meade B, Wang X (2009) Antiphonal call timing in marmosets is behaviorally significant: interactive playback experiments. J Comp Physiol A Neuroethol Sens Neural Behav Physiol 195:783–789 Available at: https://pubmed.ncbi.nlm.nih.gov/19597736/ [Accessed February 14, 2023].

Miller CT, Mandel K, Wang X (2010) The communicative content of the common marmoset phee call during antiphonal calling. Am J Primatol 72:974–980 Available at: https://pubmed.ncbi.nlm.nih.gov/20549761/ [Accessed September 1, 2024].

Miller CT, Thomas AW, Nummela SU, de la Mothe LA (2015) Responses of primate frontal cortex neurons during natural vocal communication. J Neurophysiol 114:1158–1171 Available at: https://journals.physiology.org/doi/abs/10.1152/jn.01003.2014 [Accessed December 26, 2021].

Miller CT, Wang X (2006) Sensory-motor interactions modulate a primate vocal behavior: Antiphonal calling in common marmosets. J Comp Physiol A Neuroethol Sens Neural Behav Physiol 192:27–38.

Miller CT, Wren Thomas A (2012) Individual recognition during bouts of antiphonal calling in common marmosets. J Comp Physiol A Neuroethol Sens Neural Behav Physiol 198:337– 346 Available at: https://pubmed.ncbi.nlm.nih.gov/22277952/ [Accessed February 11, 2025].

Müller-Preuss P, Jürgens U (1976) Projections from the “cingular” vocalization area in the squirrel monkey. Brain Res 103:29–43.

Nummela SU, Jovanovic V, De La Mothe L, Miller CT (2017) Social Context-Dependent Activity in Marmoset Frontal Cortex Populations during Natural Conversations. J Neurosci 37:7036–7047 Available at: https://pubmed.ncbi.nlm.nih.gov/28630255/ [Accessed December 19, 2022].

Ochsner KN, Beer JS, Robertson ER, Cooper JC, Gabrieli JDE, Kihsltrom JF, D’Esposito M (2005) The neural correlates of direct and reflected self-knowledge. Neuroimage 28:797– 814 Available at: https://pubmed.ncbi.nlm.nih.gov/16290016/ [Accessed February 11, 2025].

Pachitariu M, Steinmetz N, Kadir S, Carandini M, Harris K (n.d.) Fast and accurate spike sorting of high-channel count probes with KiloSort.

Pasternak T, Greenlee MW (2005) Working memory in primate sensory systems. Nat Rev Neurosci 6:97–107 Available at: https://pubmed.ncbi.nlm.nih.gov/15654324/ [Accessed February 11, 2025].

Paus T (2001) Primate anterior cingulate cortex: Where motor control, drive and cognition interface. Nature Reviews Neuroscience 2001 2:6 2:417–424 Available at: https://www.nature.com/articles/35077500 [Accessed January 31, 2025].

Paxinos & Watson & Petrides & Rosa & Tokuno (2012) The Marmoset Brain in Stereotaxic Coordinates, 1st Edition. Elsevier Available at: https://www.goodreads.com/work/best_book/17487503-the-marmoset-brain-in-stereotaxic-coordinates [Accessed November 8, 2021].

Roy S, Miller CT, Gottsch D, Wang X (2011) Vocal control by the common marmoset in the presence of interfering noise. J Exp Biol 214:3619–3629 Available at: https://pubmed.ncbi.nlm.nih.gov/21993791/ [Accessed February 11, 2025].

Roy S, Zhao L, Wang X (2016) Distinct neural activities in premotor cortex during natural vocal behaviors in a new world primate, the common marmoset (Callithrix jacchus). Journal of Neuroscience 36:12169–12179.

Rudebeck PH, Buckley MJ, Walton ME, Rushworth MFS (2006) A role for the macaque anterior cingulate gyrus in social valuation. Science 313:1310–1312 Available at: https://pubmed.ncbi.nlm.nih.gov/16946075/ [Accessed February 11, 2025].

Sallet J, Mars RB, Noonan MP, Andersson JL, O’Reilly JX, Jbabdi S, Croxson PL, Jenkinson M, Miller KL, Rushworth MFS (2011) Social network size affects neural circuits in macaques. Science 334:697–700 Available at: https://pubmed.ncbi.nlm.nih.gov/22053054/ [Accessed February 11, 2025].

Schaeffer DJ, Gilbert KM, Hori Y, Gati JS, Menon RS, Everling S (2019) Integrated radiofrequency array and animal holder design for minimizing head motion during awake marmoset functional magnetic resonance imaging. Neuroimage 193:126–138.

Selvanayagam J, Johnston KD, Everling S (2024) Laminar Dynamics of Target Selection in the Posterior Parietal Cortex of the Common Marmoset. J Neurosci 44 Available at: https://pubmed.ncbi.nlm.nih.gov/38627088/ [Accessed August 14, 2024].

Shadlen MN, Kiani R (2013) Decision making as a window on cognition. Neuron 80:791–806 Available at: https://pubmed.ncbi.nlm.nih.gov/24183028/ [Accessed February 11, 2025].

Shergill SS, White TP, Joyce DW, Bays PM, Wolpert DM, Frith CD (2013) Modulation of somatosensory processing by action. Neuroimage 70:356–362 Available at: https://pubmed.ncbi.nlm.nih.gov/23277112/ [Accessed February 11, 2025].

Sutton D, Larson C, Lindeman RC (1974) Neocortical and limbic lesion effects on primate phonation. Brain Res 71:61–75 Available at: https://pubmed.ncbi.nlm.nih.gov/4206919/ [Accessed January 28, 2025].

Takahashi DY, Narayanan DZ, Ghazanfar AA (2013) Coupled Oscillator Dynamics of Vocal Turn-Taking in Monkeys. Current Biology 23:2162–2168.

Teller DY (1984) Linking propositions. Vision Res 24:1233–1246 Available at: https://pubmed.ncbi.nlm.nih.gov/6395480/ [Accessed February 11, 2025].

Tian B, Reser D, Durham A, Kustov A, Rauschecker JP (2001) Functional specialization in rhesus monkey auditory cortex. Science 292:290–293 Available at: https://pubmed.ncbi.nlm.nih.gov/11303104/ [Accessed July 23, 2023].

Van Mao C, Araujo MFP, Nishimaru H, Matsumoto J, Tran AH, Hori E, Ono T, Nishijo H (2017) Pregenual Anterior Cingulate Gyrus Involvement in Spontaneous Social Interactions in Primates-Evidence from Behavioral, Pharmacological, Neuropsychiatric, and Neurophysiological Findings. Front Neurosci 11 Available at: https://pubmed.ncbi.nlm.nih.gov/28203143/ [Accessed February 11, 2025].

Walker JD, Pirschel F, Sundiang M, Niekrasz M, MacLean JN, Hatsopoulos NG (2021) Chronic wireless neural population recordings with common marmosets. Cell Rep 36 Available at: https://pubmed.ncbi.nlm.nih.gov/34260919/ [Accessed February 11, 2025].

Wallis CU, Cardinal RN, Alexander L, Roberts AC, Clarke HF (2017) Opposing roles of primate areas 25 and 32 and their putative rodent homologs in the regulation of negative emotion. Proc Natl Acad Sci U S A 114:E4075–E4085 Available at: https://pubmed.ncbi.nlm.nih.gov/28461477/ [Accessed February 11, 2025].

Wallis CU, Cockcroft GJ, Cardinal RN, Roberts AC, Clarke HF (2019) Hippocampal Interaction With Area 25, but not Area 32, Regulates Marmoset Approach-Avoidance Behavior. Cereb Cortex 29:4818–4830 Available at: https://pubmed.ncbi.nlm.nih.gov/30796800/ [Accessed February 11, 2025].

Wong RK, Selvanayagam J, Johnston K, Everling S (2024) Functional specialization and distributed processing across marmoset lateral prefrontal subregions. Cereb Cortex 34 Available at: https://pubmed.ncbi.nlm.nih.gov/39390711/ [Accessed January 21, 2025].

Wong RK, Selvanayagam J, Johnston KD, Everling S (2023) Delay-related activity in marmoset prefrontal cortex. Cereb Cortex 33:3523–3537 Available at: https://pubmed.ncbi.nlm.nih.gov/35945687/ [Accessed February 11, 2025].

Zhao L, Wang X (2023) Frontal cortex activity during the production of diverse social communication calls in marmoset monkeys. Nat Commun 14 Available at: https://pubmed.ncbi.nlm.nih.gov/37857618/ [Accessed February 11, 2025].

